# Wastelands or Preferred-lands? Indicators for redefining Desert conservation

**DOI:** 10.1101/2024.01.16.575797

**Authors:** Manasi Mukherjee, Dhriti Banerjee, Indu Sharma, Mitali Mukerji

**Affiliations:** Jodhpur City Knowledge and Innovation Foundation, Indian Institute of Technology, Jodhpur, Rajasthan, India; Desert Regional Centre,, Zoological Survey of India, Jodhpur, Rajasthan, India; Department of Bioscience and Bioengineering, Indian Institute of Technology, Jodhpur, Rajasthan, India

**Keywords:** Ecoregion, Thar Desert, Spatio-temporal variation, Avian Diversity, Species richness, feeding behaviour

## Abstract

Desert ecosystems though crucial in shaping global climate, are often perceived to have limited function and are frequently designated as wastelands due to low biotic diversity and abundance. We explore the functional properties of desert through assessment of spatio-temporal variations of avian species richness and insect abundance of the relatively understudied Thar in Rajasthan. The study was conducted across four contrasting ecoregions (Western Thar (WT), Eastern Thar (ET), Transitional Zone (TZ), and Cultivated Zone (CZ) spanning 33 counties of Rajasthan. Monthly avian diversity were obtained from crowdsourced eBird data. Insect diversity and abundance accross the ecoregions were curated from the annual surveys of ZSI, India. The leads were corroborated with five years feeding behaviour observational data recorded from a representative site spanning 852 acres. There was a significant seasonal differences in bird species richness between the ecoregions with the most pronounced variations between summer and winter and CZ exhibiting the highest spatial variability followed by WT. WT also recorded the highest insect diversity and abundance, especially the orders Orthoptera, Hymenoptera, Coleoptera, Lepidotera and Diptera. The correlation matrix between bird and insect abundance and Kernel Density Estimation (KDE) plot indicated the relation between diverse dietary preferences and migratory patterns. These observations challenge the conventional notion that resource-limited environments harbour lesser biodiversity especially in arid environments. The dominance of insectivory among winter migrants aligns with the higher density of resident arthropods and emphasizes the pivotal role of insects as a food source during the migratory period. The adaptability of resident species to varied food sources, particularly arthropods, provides a means to strategically navigate during resource constraints. This study thus provides a unique framework to (a) redefine the functional importance of desert using migratory birds as prominent ecological indicators. (b) include an ecological criteria during policy formulation for biodiversity conservation. It also emphasizes reevaluation of deserts during designation of wastelands especially in the unique WT eco-regions when evolving developmental as well as ecological restoration strategies.

## 1. Introduction

Deserts, often perceived as barren ecosystem, on closer examination of the trophic structures reveals a rich biodiversty. In contrast to tropical biomes (Jetz et al., 2012), resource-limited deserts pose a challenge for sustenance of all biota (She et al., 2018). Desert biomes face challenges like high temperature fluctuations and limited resources, intensifying competition among habitants, both residents as well as migrants (Zimin et al., 2023). Resident birds evolve adaptive strategies to mitigate this competition that includes habitat and temporal niche partitioning, dietary flexibility, and niche specialization (Higginson, 2017) For migratory birds, in addition to favorable temperatures and environmental conditions, the availability of sufficient food and nesting sites (Carey, 2009) plays a crucial role in selecting migration destinations.

The Thar desert spanning 200000 sq kms is a vast arid region in the north western part of India and is a host to 682 species of flora, and 1195 (2.12 % of Indian fauna) species of fauna (Roy and Roy, 2019). As the world’s most densely populated desert, the Thar desert contends with on-going human-induced pressures, leading to various adaptations to support human and faunal habitation (ul Islam and Rahmani, 2011). Understanding why migrating birds return to resource-limited biomes, like deserts, is crucial for shaping conservation strategies, especially considering the vulnerability of these environments to climate and developmental changes. In the present study we explore the functional properties of Thar desert through assessment of spatio-temporal variations of avian species richness and insect diversity and abundance accross four delineated ecoregions of Thar from our earlier study (Mukherjee et al., 2023). Briefly these include, Western Thar (WT), Easter Thar (ET), Transitional Zone (TZ) and Cultivated ZOne (CZ).

The crowdsourced data from ebird and survey data from ZSI, accross 33 districts/ counties of Rajasthan was curated and analysed for the study. The bird species richness data in Thar’s ecoregions revealed seasonal variation, indicating an influx of migratory species. The WT, that is the most arid part of Thar stood out as a preferred region for winter migrants, with insects identified as a major food source. Resident birds adapt to ephemeral resources, while migrants rely on specific food sources, emphasizing the importance of maintaining ecological balance.

This study for the first time underscored the need for carrying out a functional ecology of deserts prior to designating them as wastelands for developmental activities. Also, it recognizes birds and insects as indicators of ecologically sustainable ecosystems.

## 2. Methods

We investigated the significance of biota in shaping the functional characteristics of the desert by examining the following two hypotheses.

1. Significant seasonal difference in species richness of birds between four ecoregions of Thar
2. Insects are the major source of food causing winter migration of birds in Western Thar

### 2.1. Study area

Thar in Rajasthan situated between 27.4695° N to 70.6217° E (1) features the Aravallis, India’s oldest mountain range, that geographically divides the region into Western Thar (WT), comprising 80% of the Thar desert, and the relatively greener Eastern Thar (ET). Thirty three districts of Rajasthan were selected to analyze the differences in avian species richness in each ecoregion Mukherjee et al. (2023) and their relation to the Insecta of Thar. The findings were validated through the feeding behaviour observational data obtained from the selected sub-site (Indian Institute of Technology-Jodhpur (IITJ) campus) situated in Jodhpur, a district of Western Thar.

Bird diversity and their behavioural ecology were studied from this selected sub-site during September 2019 to September 2023. The campus spanning an area of 852 acres provides an amalgamation of both undisturbed serene desert grassland environment and human settlements like buildings for academics, laboratories, administration, health care and residences all guarded by large berms to withstand strong sandstorms (Fig. 1.

**Figure 1:**
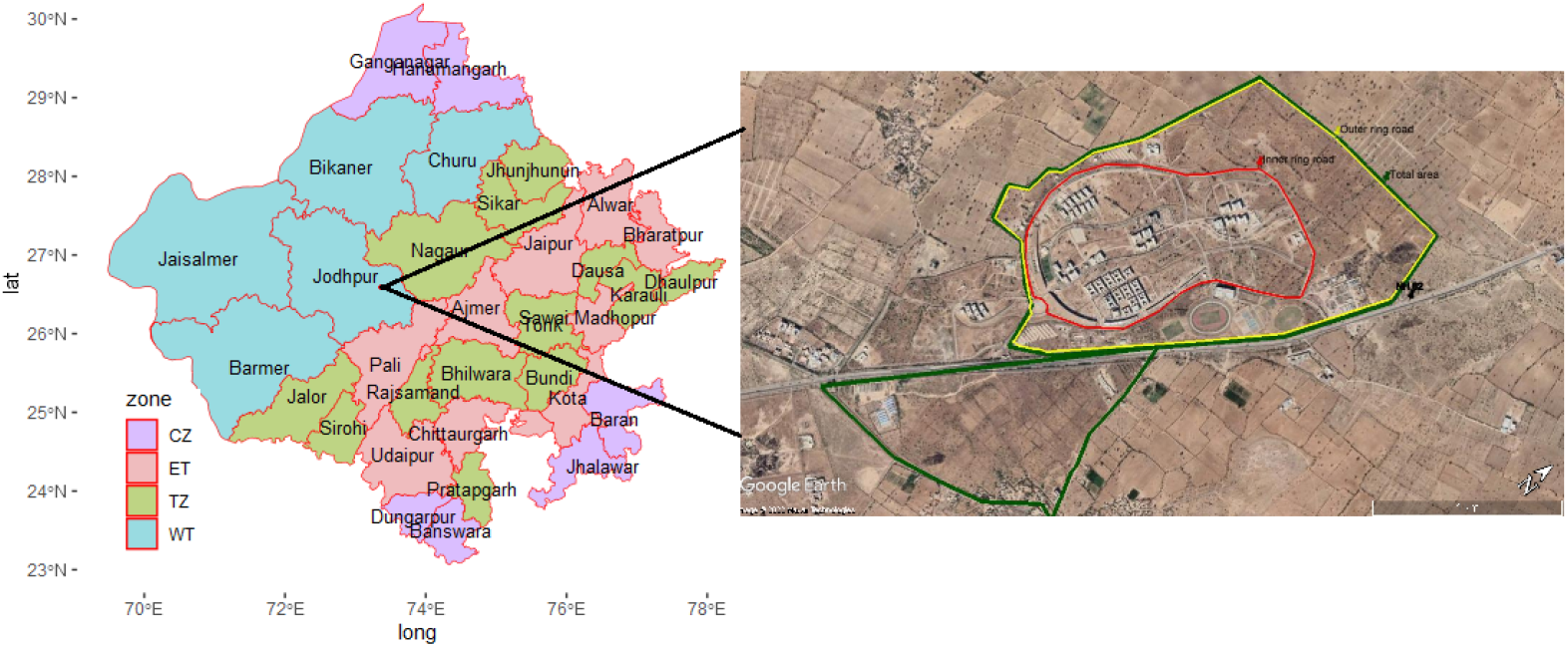
(a) Sample site: A map of Rajasthan delineating districts, portraying biotic-ecoregions using color codes; Western Thar (WT) represented by blue, Eastern Thar (ET) by pink, Transitional Zone (TZ) by green, and Cultivated Zone (CZ) by purple. (b) Validation site: The WT representative site is situated within IIT Jodhpur spanning 852 acres. Arid vegetation predominantly covers the outer boundary of the campus, denoted by green lines, while red lines outline the inner boundary enclosing the primary constructed area. The regularly covered area for data collection is highlighted by a yellow line.

### 2.2. Data collection and analysis

#### 2.2.1. Sample data: Aves

Avian species richness data from all four ecoregions of Rajasthan were obtained from eBird. This was curated from the histogram data under the barchart category available in eBird (https://ebird.org/barchart?r=IN-RJ&yr=all&m=) as a matrix of columns. All the districts under each ecoregions from our earlier study Mukherjee et al. (2023) namely Western Thar (WT), Eastern Thar (ET), Transitional ZOne (TZ) and Cultivated Zone (CZ) were selected to curate the species richness (S) data.

#### 2.2.2. Spatio-temporal variation analysis of avian diversity

Ecoregion-wise mean seasonal abundance was calculated for analysing temporal variations. A Student’s t-test on species richness between ecoregions were performed to analyze the spatial differences. And the corresponding results were represented in a matrix of p-values (cut off *≤* 0.05).

Percentage difference of species richness was calculated between each season (Winter, Summer and Monsoon) for all the ecoregions (WT, ET, TZ and CZ). Months from October to February were considered winter, March to June as summer and July to September as monsoon.

#### 2.2.3. Sample data: Insecta

Order level insecta diversity and abundance were studied in Rajasthan between 2017 and 2022 through systematic random sampling conducted across all districts by Zoological Survey of India (ZSI). Insect collection involved various techniques such as employing Aerial, Aquatic, and Sweeping Nets. Aerial nets were specifically utilized for capturing butterflies and other flying insects, while the Aquatic net facilitated the collection of insects from water bodies and Sweeping nets for terrestrial insects. Additionally, a Light trap was strategically used to attract and collect various insect orders drawn to light during the night.

Post-collection procedures involved placing the gathered insects in a Killing Bottle containing Ethyl acetate for preservation. Immediate mounting of collected specimens took place, with certain insects being relaxed before mounting to prevent them from becoming excessively hard and brittle over time.Relaxation was achieved by placing the insects in a wide-mouth, air-tight jar filled with moist sand and covered with blotting paper. A few drops of carbolic acid were added to prevent mold formation during this process, lasting one to two days. Subsequently, pinning of the specimens was carried out, and the pinned insects were meticulously arranged on a setting board before being left to dry in a drying chamber.

Specimens were then labelled with scientific classification, locality, host plant, date of collection, and the name of the collector. An aspirator played a role in collecting small and soft-bodied insects. Following the collection, a preservation solution of 70-75% alcohol with a small quantity of glycerine was added to the specimen tube. Additionally, very small insects were preserved by mounting them on permanent slides.

The collected data was binned into the same ecoregions described above to analyze the distribution of insecta across Rajasthan. Order-wise abundance variation of these insects in each ecoregion was analysed. Additionally, this data was employed to explore potential relationships between the bird and insect populations of these ecoregions.

#### 2.2.4. Establishing the study area for validation

The validation study area, encompassing around 852 acres, was chosen within the Jodhpur district of Western Thar (WT), situated on the IITJ campus. Feeding behavior study of birds was carried out in this designated validation site, consisting of 66% of desert shrubland and grassland, and 34% of constructed areas. Alternate day observations were made during early morning (6:00-7:00 am) and early evenings (5:00 pm) hours. Crepuscular birds were mostly viewed towards the beginning or end of the trail. Due to the unavailability of necessary tools, nocturnal birds were not covered in the study. The time of observations varied based on seasons, sunrise and sunset. The buildings, berms, pipe ways, open drainages, gardens, playgrounds, dump yards and parking areas were also regularly observed. Change in trails were often made to ensure maximum coverage of the study area and avoiding missing of any sightings. Lamp posts, streetlights and tip of the berms were kept in watch for perching activity of birds like raptors and other carnivores. Fruiting season of major plants like Neem, Plums, Ker, Babul and Khejri were noted to observe the feeding efficacy of foragers.

#### 2.2.5. Validation data for analyzing feeding behaviour

The avian species recorded were identified following standard literature Raju and Ramachandran (2017); Vyas (2013) and websites like eBird and Aviase. Photos and videos were captured using Panasonic handicam HC-V180. Photos of all the species recorded are uploaded on the website of the author under the link <https://manasimukherjee.info/2021/05/22/birds-of-iit-jodhpur/> for reference. Based on their appearance in the campus, the recorded species were classified into Residents, Summer migrants, Winter migrants and Monsoon migrants.

Throughout the study duration, variations in food and feeding behaviours of the identified bird species were documented across seasons. The available food options for campus birds were classified into grains, fruits, nectar, insects, and vertebrates. Interrelation between the classified birds and their food types was assessed through a correlation matrix. To assess food preferences, each classified food type was assigned a score out of 10 per bird species for each season. Using this scoring system, a kernel density plot was created to examine seasonal preferences between migrant and resident bird populations on the campus.

## 3.. Results

The data collected on bird diversity in the study area (IIT Jodhpur campus) from 2019 to 2023 revealed a notable increase in avian presence during the winter months compared to other seasons. Given that multiple factors could influence winter bird migrations, we explored the significant determinants that could influence the migration of birds in WT through

a. Spatio-temporal variation analysis of avian diversity across four ecoregions
b. Validation through comparative assessment of major food sources of migrants and resident avian species of WT.

### 3.1. Significant seasonal differences in species richness of birds between four ecoregions of Thar

The Species richness (S) obtained from ebird data on 33 counties of Rajasthan state of India accross four ecoregions (Mukherjee et al., 2023) is illustrated in Table 1. The seasinal species richness of birds varied significantly between the ecoregions (Fig. 2). The richness of the birds remained highest during winters followed by monsoons and was least during summers in all the ecoregions.

**Table 1:** Seasonal variation in Species richness (S) of birds in each ecoregion of Thar.

| Seasons | Months | WT | ET | TZ | CZ |
| --- | --- | --- | --- | --- | --- |
| **Winter** | **October** | 297 | 357 | 299 | 205 |
|  | **November** | 306 | 367 | 311 | 193 |
|  | **December** | 304 | 378 | 317 | 170 |
|  | **January** | 293 | 389 | 319 | 181 |
|  | **February** | 292 | 390 | 329 | 216 |
| **Summer** | **March** | 252 | 376 | 302 | 145 |
|  | **April** | 210 | 337 | 267 | 76 |
|  | **May** | 183 | 286 | 229 | 78 |
|  | **June** | 167 | 265 | 208 | 81 |
| **Monsoon** | **July** | 191 | 260 | 209 | 100 |
|  | **August** | 239 | 289 | 247 | 94 |
|  | **September** | 283 | 326 | 286 | 91 |

**Figure 2:**
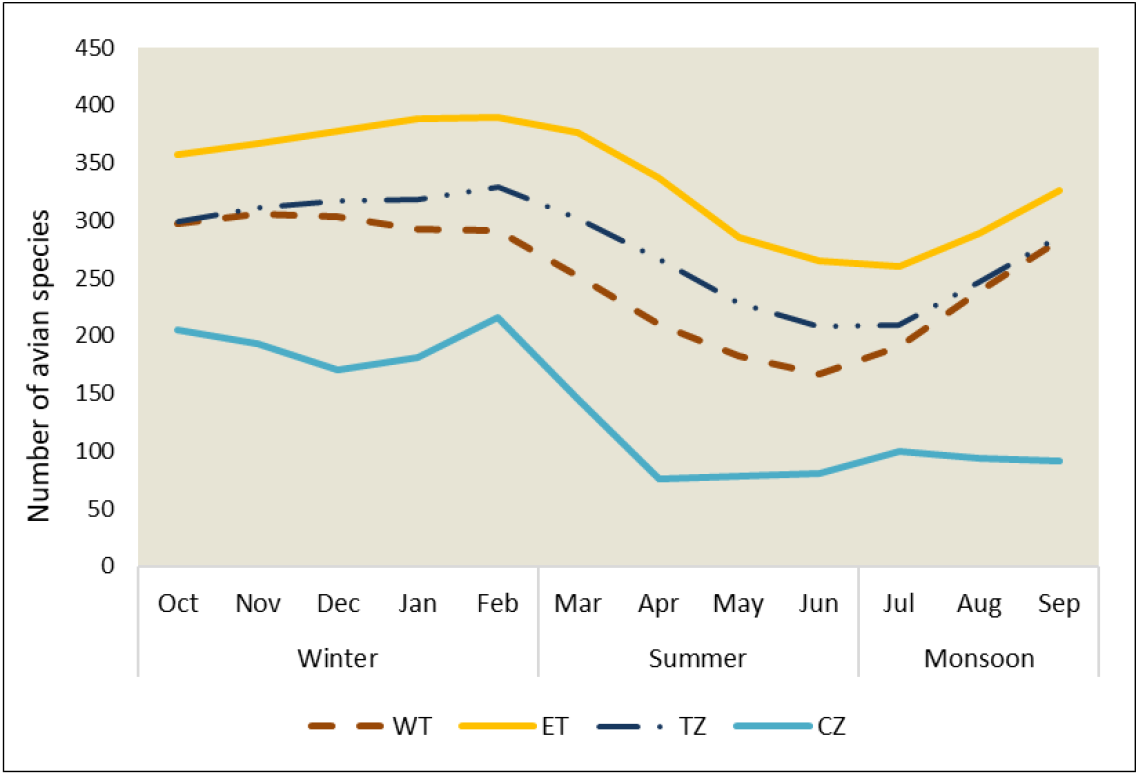
Spatio-temporal variation in avian species richness (S) across four ecoregions within Thar. The species richness of birds in four ecoregions of Thar exhibited monthly variations. Throughout the year, Eastern Thar (ET) consistently displayed the highest species richness, followed by Transitional Zone (TZ), Western Thar (WT), and Cultivated Zone (CZ). Seasonally, species richness remained at its peak during winter across all ecoregions, whereas it was at its lowest during summer.

The p-value matrix (Table 2) showed that the monthly variation in birds species richness of all the four ecoregions differed significantly from each other. The only exception remained between Western Thar and Transitional Zone, where the differences are non-significant. The percentage of seasonal difference in bird species richness indicated the following

**Table 2:** p-value matrix of seasonal avian species richness (S) difference between each ecoregion.

|  | WT | ET | TZ | CZ |
| --- | --- | --- | --- | --- |
| WT | 1 | 0.00051 | 0.2067 | 2.45E-05 |
| ET | 0.0005098 | 1 | 0.005635 | 3.89E-09 |
| TZ | 0.2067 | 0.005635 | 1 | 6.71E-07 |
| CZ | 2.45E-05 | 3.89E-09 | 6.71E-07 | 1 |

- Temporally, the highest difference in species richness occurs between summer and winter across all ecoregions
- Spatially CZ shows highest variation in seasonal bird species richness, followed by WT, ET and TZ

Futhermore, Figure 3 also indicates that WT has comparatively larger percentage of winter migrants as compared to ET or TZ.

**Figure 3:**
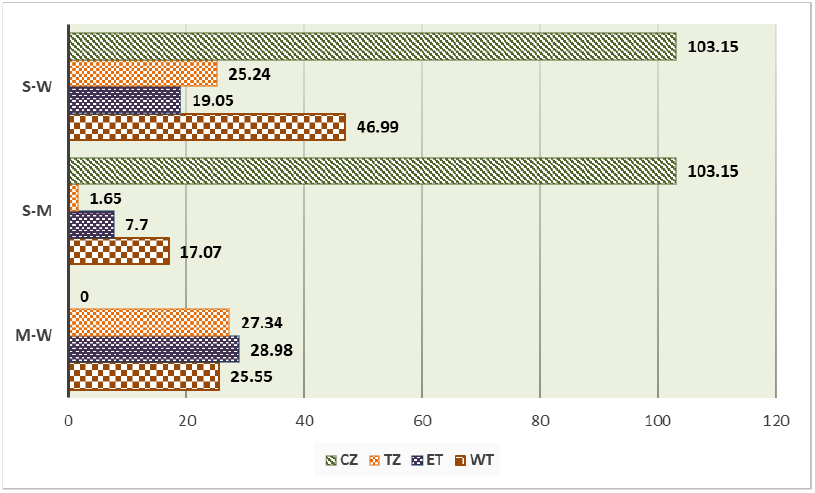
Percentage variance in species richness between seasons among the four ecoregions of Thar: Western Thar (WT), Eastern Thar (ET), Transitional Zone (TZ) and Cultivated Zone (CZ). ‘S’ denotes ‘Summer’, ‘W’ as ‘Winter’, and ‘M’ as ‘Monsoon’. The percentage differences between seasons are indicated at the apex of the bars. CZ exhibited the greatest percentage difference between summer and winter, followed by WT, ET, and TZ.

### 3.2. Seasonal patterns in avian diversity of a longitudinal cohort representing WT

Due to the severe aridity and limited resources in the Western Thar region, the higher abundance of migratory birds compared to other ecoregions seemed paradoxical. Another important question that emerges is, considering the scarcity of substantial vegetation to sustain them, “what resources cause this higher percentage of avian winter migration in WT ?” To address the above, an ongoing survey data which spans from 2018 to 2023 was used from selected representative site of WT for analyzing the bird diversity and their feeding behaviour. The diversity and behavioural data collected show the presence of 80 avian species encompassing 10 taxonomic orders and 39 distinct families (Table 3). Among the catalogued avifauna, 43 species were identified as residents, 19 as winter migrants, 10 as monsoon and 8 as summer migrants. The campus exhibited pronounced seasonal fluctuations in bird diversity. Prominent among the winter migrants were species such as rosy starlings (*Pastor rosseus*), Siberian stonechats (*Saxicola maurus maurus*), black redstarts (*Phoenicurus ochruros*), Eurasian wrynecks (*Jynx torquilla*), and variable wheaters (*Oenanthe picata*). Siberian stonechats displayed limited presence, often appearing in pairs. Noteworthy was presence of the subspecies of the Eurasian Wryneck, *Jynx torquilla, himalayana* in the area. Variable wheater with two visible morphs: picata and capistrata and black redstart, two other winter migrants, briefly inhabited the campus. Several shrike species frequented the campus, with bay-backed, long-tailed, and great-grey shrikes emerging as the most prevalent. Their numbers dwindled during winter and surged from summer to the monsoon season. The elusive Philippine brown shrike makes a rare appearance during winter, known for its preference to feed on larger insects like wasps and grasshoppers. In addition to the winter migrants, the campus also hosted monsoon migrants like baya weaver (*Ploceus philippinus*). The green bee-eater (*Merops orientalis*), another avian visitor, engaged in local migration, settling within the campus during late summers. Blue-cheeked bee-eater (*Merops persicus*), another migratory species, resided in the campus for the shortest interval (June to September) in limited numbers, potentially making a brief stopover.

**Table 3:**
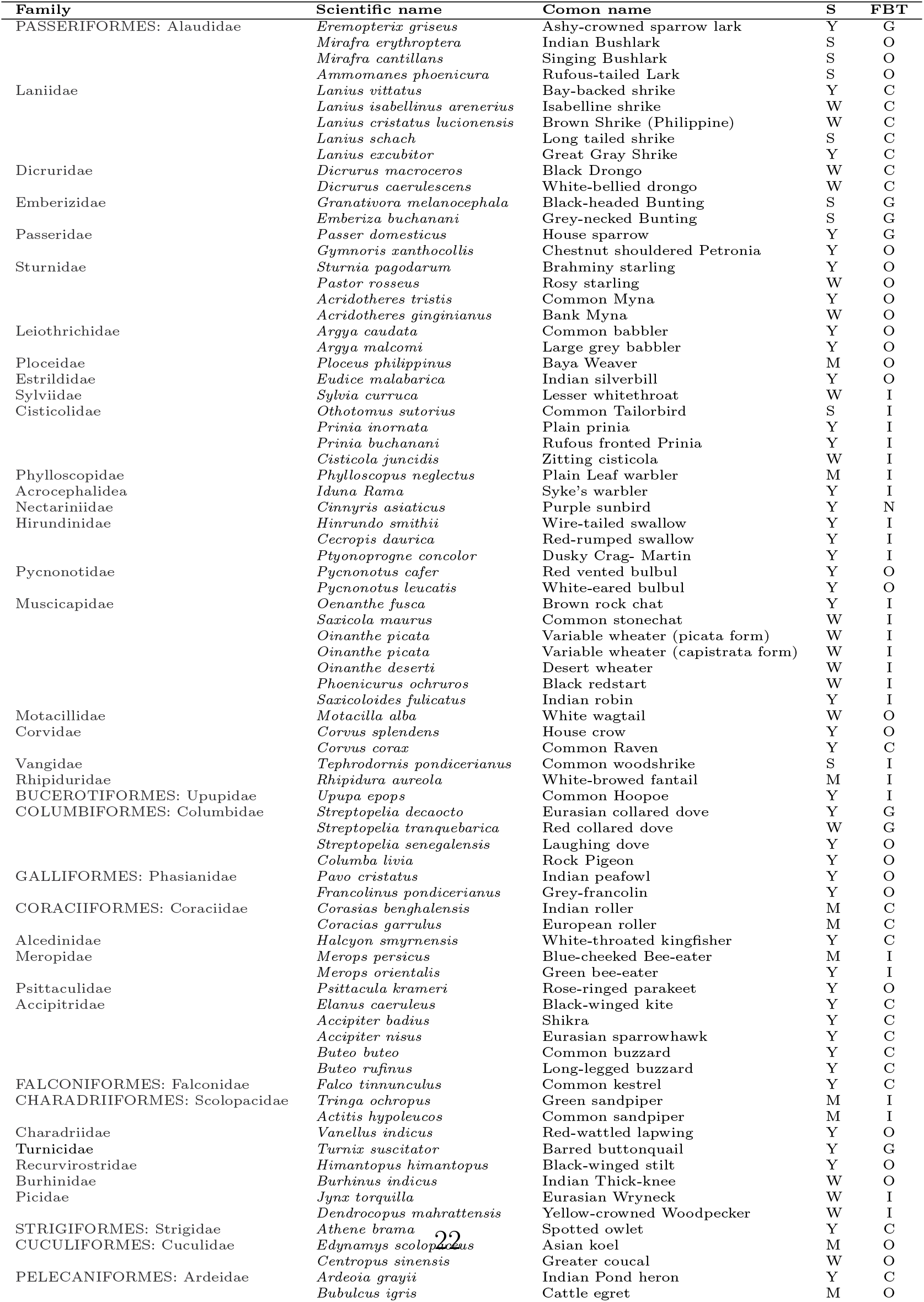
List of bird species recorded, their seasonal appearance (S) and Feeding Behaviour Type (FBT) from IITJ campus.

The surge of the listed bird species during winter on the campus prompted a need to understand their feeding behaviours. Further observations and analysis were sought to grasp the primary reasons driving their arrival in an ecosystem constrained by limited resources. Consequently, this exploration led to the formulation of the second hypothesis.

### 3.3. Bird’s dietary preferences in WT

Based on the observations of feeding behavior conducted during the study period, the birds were categorized into distinct groups of granivores (consumers of seeds), frugivores (consumers of fruits), nectarivores (consumers of nectar), insectivores (consumers of insects, including other arthropods), and carnivores (predators of both small and large vertebrates). Noteworthy, due to the extreme climatic conditions, a substantial proportion of the bird population were observed to have developed omnivorous and opportunistic feeding tendencies (Table 3). The highest proportion were omnivores (33.75%), followed by insectivores (32.5%), carnivores (23.75%), granivores (8.75%), and nectarivores (1.25%). Despite the significance of fruit consumption within the omnivorous feeding behavior, it’s worth noting that none of the observed birds displayed a purely frugivorous nature. To evaluate the relationship between the seasonal migration patterns of bird species in the research area and their feeding habits, a correlation matrix (Table 2) was generated based on the gathered data. The correlation matrix showed a significant negative correlation (r= -0.6) between winter migrants and residents with respect to feeding behaviours and between insectivores and granivores (r= -0.53). The matrix did not yield any other notably high positive (r*≥*0.5) or negative (r*≥*-0.5) correlation coefficient values. The matrix contributed to a partial comprehension of the matter by elucidating the varying significance of food sources within each migratory category. Among the resident bird species, there was a stronger inclination towards carnivory, nectarivory, and frugivory. During the summer migration period, a preference for granivory was evident, whereas winter and monsoon migrants displayed a predilection for insectivory and frugivory.

The seasonal preference of food type among migrants of the studied representative site was corroborated by Kernel Density Estimation (KDE) plot in Fig.5. Remarkably, resident species seem to exhibit more familiarity and awareness of the available food types within the ecosystem, that predominantly manifests in their omnivorous nature.

As evident from the probability curve (Fig. 5), although the density of arthropods and vertebrates is relatively low among the resident food sources, their consumption frequency remains higher compared to other food types. This signifies that the resident species exhibit heightened adaptability to the constraints of limited resources in the ecosystem. A conspicuous revelation is the consistent requirement for arthropods in the dietary habits of both residents and migrants. Conversely, during the summer migration period, there is a lower density but a higher frequency of consumption towards grains and arthropods among the migrants. Monsoon migrants, in contrast, strike a balance in their feeding behaviour with a greater density of vertebrates and an elevated frequency of arthropod consumption. Winter migrants exhibit a higher density across all food types, with a pronounced frequency of consumption towards insectivory.

### 3.4. Insect distribution and abundance contribute to spatio-temporal differences in avian diversity across ecoregions

To verify if the findings from the representative cohort were applicable across the entire ecoregion, the study compared insect abundance in WT with that in other ecoregions. The insect data obtained from different ecoregions of Thar (Rajasthan) (Fig.6a and 6b), also indicated that WT has the highest abundance of Insecta followed by ET, TZ and CZ respectively. This corroborates the previous results, wherein the higher abundance of insecta in WT ecoregion provides ample food resource for the winter migrants that show higher affinity to insecta.

A detailed look into the order-wise abundance of insects in all the four ecoregions of Thar shows a notable difference in community composition. WT has a notably high abundance of the orders Orthoptera, Hymenoptera, Coleoptera, Lepidotera and Diptera (Fig. 7) when compared to other ecoregions of Thar.

## 4.. Discussion

Deserts, typically seen as ecosystems with restricted biodiversity and limited functionality, are frequently considered as barren lands that can be used for developmental initiatives. This study provides a unique framework for integration of behavioural ecology for comprehensive developmental activities especially in the desert ecoregions of Thar. Our study reveal that the spatial distribution and community structure of migratory birds in Thar are intricately linked to their food preferences and available resources. The most arid regions ”WT” hosts a greater diversity of bird species more prominently during winter compared to lusher environments. Also in contrast to other ecoregions within Thar, this ecoregion experiences a greater influx of avian species during winter. The driving force behind the winter influx of birds in WT seems to be the rich abundance of insects that serve as the primary source of food. Moreover, the predation of these insects by winter migrants contributes to the control and regulation of insect populations in Thar, effectively maintaining the trophic structure of the desert ecosystem.

The present study revealed significant seasonal differences in the species richness of birds between the four ecoregions of Thar (Fig. 2). The ecoregions, ET and TZ that remained higher in species richness are areas with higher precipitation than WT (IMD, 2022) and thus support resourceful vegetation. WT is predominantly a ‘thorn forest type’ or ‘scrub forest type’ whereas the ET and TZ have ‘dry deciduous forest’ Reddy et al. (2011). While the former provides challenging environment and limited resources for frugivores, nectarivores and granivores, the latter is comparatively abundant.

Our research also reveals spatial and temporal variations in avian diversity within ecoregions. Both zones with lower species richness (CZ and WT) display a relatively higher percentage of seasonal fluctuations in avian species richness, notably from summer to winter (Fig. 3). WT, in particular, was noted to exhibit the second-highest alpha diversity (395) concerning bird species (Mukherjee et al., 2023). This indicates abundant food resources in both these ecoregions, especially during winter, essential for sustaining migrating birds. Notably, the most significant percentage difference occurs in CZ, where, as suggested by Mukherjee et al. (2023), bird biodiversity is primarily influenced by its crop yield. Thar possesses distinctive geographical features that support varying species richness or communities in different seasons (Mathur and Sundaramoorthy, 2016; Mehra and S.P., 2008). The diverse components across Thar that have resulted in the formation of distinct biotic ecoregions (Mukherjee et al., 2023) now indicate ecoregion-based functioning of Thar.

A negative correlation between winter migrants and resident species concerning food availability, indicates a reduced competition within WT (Figure 4. Additionally, winter migrants exhibited a stronger correlation with arthropods (Fig. 4), compared to other available food categories that serve as a food source for primary and secondary consumers such as birds Ayal (2007). In addition, a negative correlation between granivory and insectivory (Fig. 4) also indicates minimal competition over limited resources like grains. The findings underscore the remarkable adaptability of the bird species in the study area, as they have adjusted their feeding habits in response to the challenging climatic conditions, resulting in a diverse array of feeding throughout the community. Nonetheless, the residents excel in their adaptability to the available food resources within the ecosystem, thus mitigating any potential resource competition or threat to their sustenance posed by the migrants. These results were corroborated by the observations that, winter migrants of WT showed comparatively higher density and observational score for feeding towards arthropods (Insecta) (Fig. 5).

**Figure 4:**
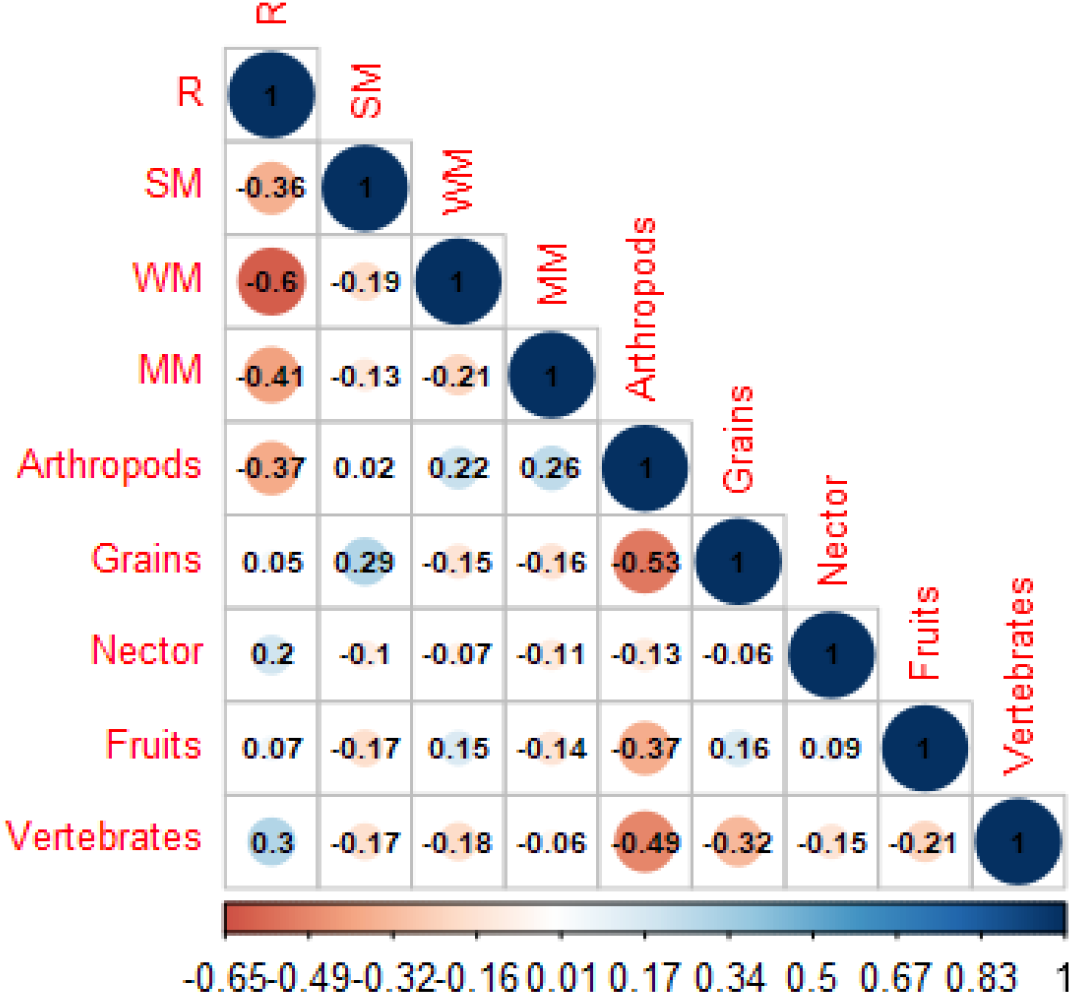
Matrix depicting the correlation between bird migration and feeding behavior at the validation site. In this matrix, ‘R’ stands for ‘Resident species’, ‘SM’ for ‘Summer migrants’, ‘WM’ for ‘Winter migrants’, and ‘MM’ for ‘Monsoon migrants’. The major classified food items of the birds observed in the study area are Arthropods, Grains, Nectar, Fruits, and Vertebrates. The color scale of the plot ranges from -1 to 1, where values closer to negative (depicted in red) signify negative relationships between migrants and specific food types, while those closer to 1 (shown in blue) indicate positive relationships. The intensity of the color shading signifies the strength of these positive or negative correlations. Size of the coloured circles is directly proportional to the size of the correlation values. A negative correlation in feeding behavior between winter migrants and resident birds suggested decreased competition within WT. Negative correlation between food types grains and arthropods and vertebrates indicate limited competition for these food types.

**Figure 5:**
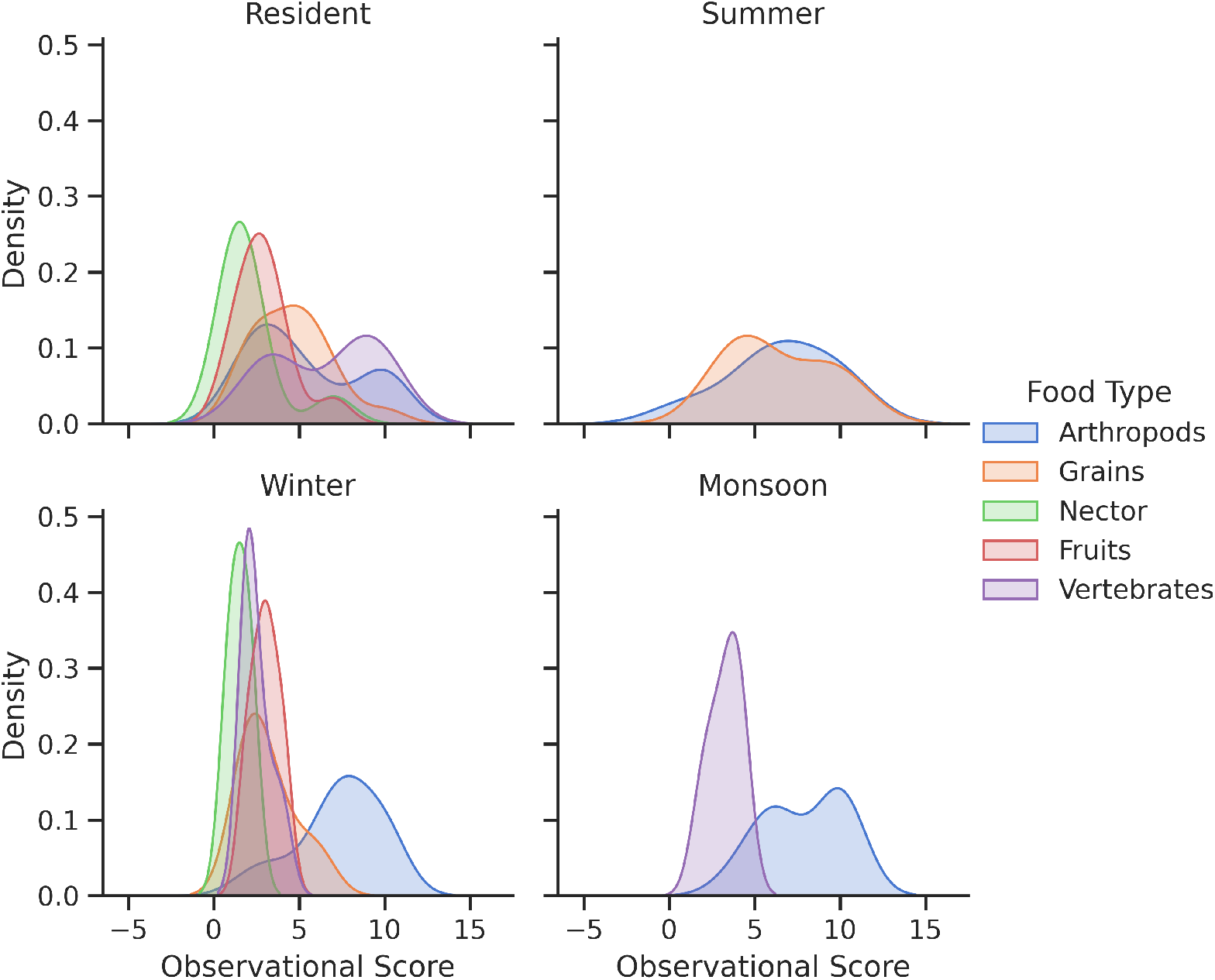
Kernel density estimate (KDE) plot for representing the feeding behaviour and seasonal appearance data using a continuous probability density curve. The x-axis represents observational scores derived from feeding behavior data at the validation site. On the y-axis, density is determined by the frequency of species and the usage of categorized food types. Residents, contrasted with all migrants, exhibit a broader utilization of various food types and display a lesser preference for arthropods, indicating an omnivorous feeding behavior. Conversely, migrants demonstrate a stronger inclination towards arthropods compared to other food types.

The spatial and temporal variations in food preferences indicate the functional significance of ecoregions, necessitating a deeper understanding of the distribution of food sources, particularly insects. Analysis performed on the distribution and abundance of insects establish their importance as a food source for migrants (Fig. 6a and 6b). The deserts or arid ecosystems are known to have low primary productivity which is sporadic in time and space Mathur and Sundaramoorthy (2016); Polis and Lubin (2005). It is possible that the detritus of the desert which decomposes very slowly due to the absence of moisture could be main source of nutrients for the detritivores. These detritivores, especially the macro-detritivores functionally generate vertical nutrient recycling loops providing nutrients to plants in dry sandy soil Sagi and Hawlena (2021). Studies have shown that Sagi et al. (2021), the macro-detritivore activities decrease during summer and increase thereafter. During summers, Western Thar (WT) exhibited limited bird species richness (Fig. 2) and resident biodiversity (Table 1). This smaller community might not sufficiently manage the high insect density in Thar (Fig. 6a and 6b). Section 3.3 also highlights the adaptable feeding behavior of resident avian populations in Thar based on seasonal resources. Conversely, migrants exhibit a preference for insects, contributing to increased avian species richness during winters and aiding in the regulation of insect populations, thus maintaining the trophic structure of the desert ecosystem.

**Figure 6:**
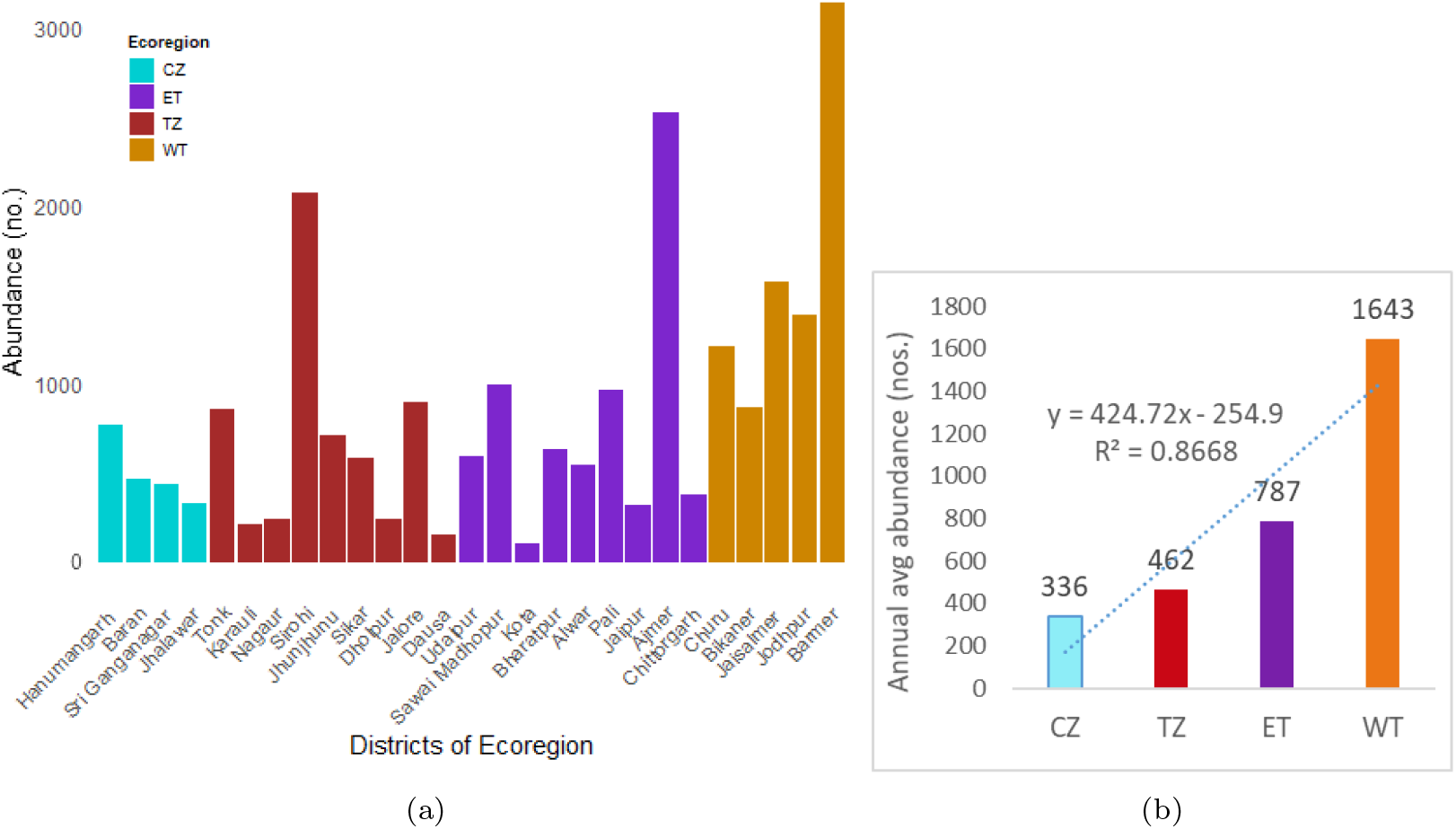
(a) Distribution of Insecta abundance across 33 districts within the four ecoregions of Thar. Among these districts, those within Western Thar (WT) exhibited the highest abundance, followed by Eastern Thar (ET), Transitional Zone (TZ), and Cultivated Zone (CZ). (b) Annual average abundance of Insecta across the four ecoregions of Thar. The average abundance value for each ecoregion is displayed at the top of the bars. The linear trendline and coefficient of determination (*R*^2^) demonstrate a notable increase in Insecta abundance from CZ to WT, indicating a significant trend.

Our observations on significantly higher insect abundance in the WT ecoregion (Fig. 6b) especially of the groups hymenoptera, coleoptera, orthoptera, and diptera (Fig. 7 corroborate with the studies from similar desert ecosystems. These groups act as the major macro-detritivores of desert ecosystems (Crawford, 1979) and comprise 80% of avian insect diets Mwansat et al. (2015). Moreover, similar to our current study, orthopterans have been documented as a year-round food source for birds in semi-arid regions of South Africa (Kok and Louw, 2000). Thus the higher abundance of these insect groups becomes a key factor (food source) to regulate the influx of winter migrants (birds) in WT. This substantiates our hypothesis that higher abundance of insecta could drive the influx of winter migrants in WT.

**Figure 7:**
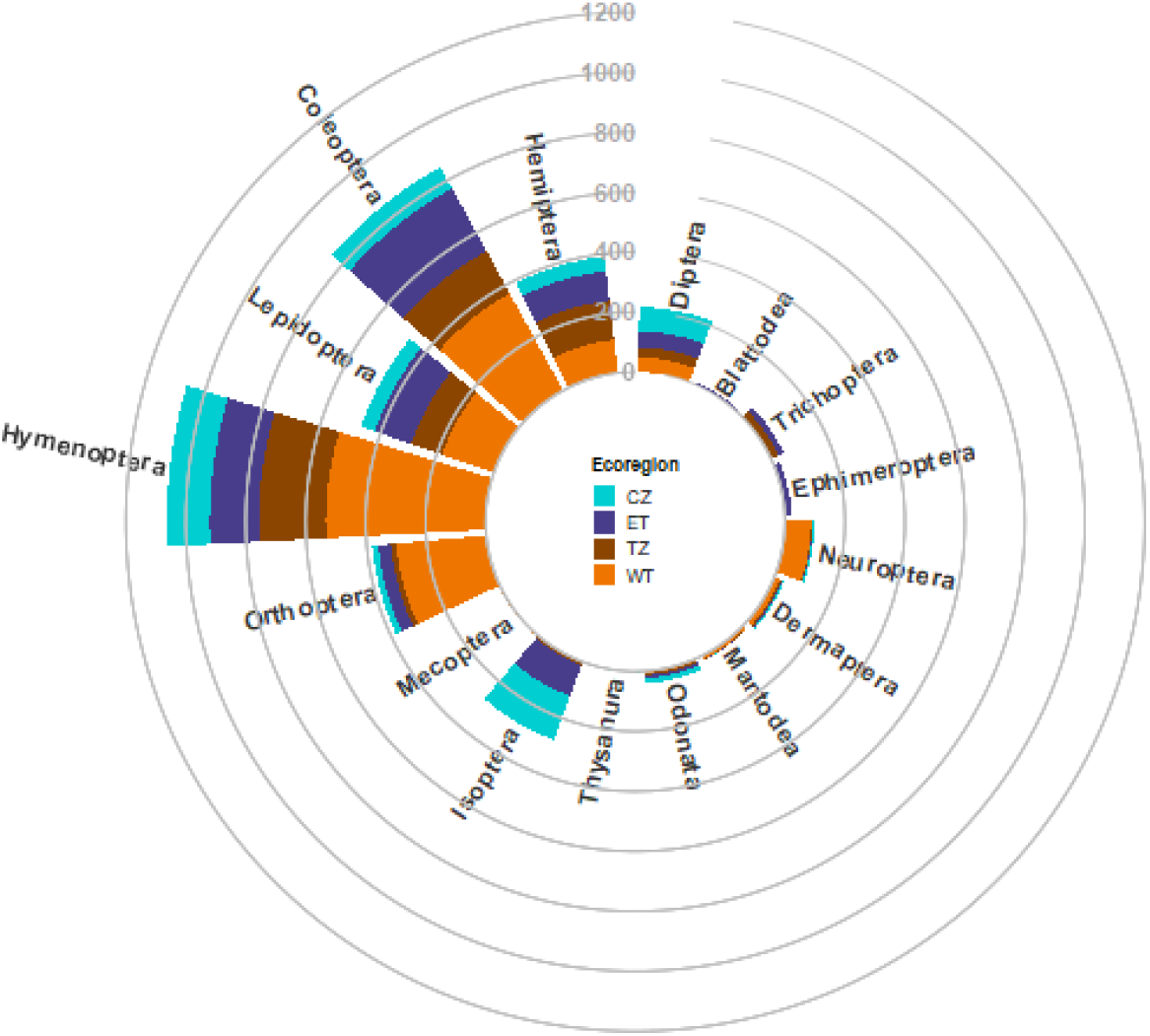
Circular bar chart displaying the spatial distribution of Insecta orders and their abundance recorded across the ecoregions of Thar. The y-axis is represented by concentric circles, indicating abundance in numbers. A survey conducted between 2017 and 2022 assessed the abundance of the Insecta class, encompassing 16 orders, within the Thar area. The stacked bars illustrate the relative abundance of insect orders in these ecoregions. While Eastern Thar (ET) and Transitional Zone (TZ) exhibited similar levels of insect abundance, Cultivated Zone (CZ) demonstrated the lowest abundance of insects. Within Western Thar (WT), a notable number of orders, particularly Hymenoptera, Orthoptera, Coleoptera, Lepidoptera, and Neuroptera, displayed comparatively higher abundance. The order Isoptera showed significant abundance in CZ and ET but notably lower abundance in WT.

About 27.38% of Rajasthan’s terrain is classified under the Sands-semistabilized-stabilized moderate high wasteland category (NRSC, 2019). Our research indicates that WT serves as a biodiversity hotspot, fostering a diverse population of migratory birds due to its abundance of insects. Therefore, our study challenges the prevailing belief in the limited functionality of limited functionality of arid ecoregions and emphasizes revisiting the characterization of these regions as wastelands, especially in WT. However, the lack of specific avian species abundance data in Thar limits understanding the impact of insect abundance on the population size of migratory birds. More comprehensive studies on the temporal and spatial variations in vegetation and other significant taxa, along with their interconnectedness, are crucial to grasp the intricate functionalities and interdependencies within ecoregions.

In conclusion, from our study WT stands out as a highly favored ecoregion for migratory birds during winters primarily driven by a higher abundance of insects in WT. This underscores the need for (a) re-assessing deserts during the categorization of wastelands, considering their resident and migratory biodiversity. (b) ecoregion-specific conservation strategies to address ongoing anthropogenic activity and climate change, preventing accelerated biodiversity loss in these vulnerable environments.

## Acknowledgements

We express our gratitude to Prof. Santanu Chaudhury, Director of IIT Jodhpur, for his consistent support throughout this endeavor. Author Manasi Mukherjee acknowledges JCKIF for providing salary support under the Thar-DESIGNS project.

## Authorship contribution statement

**Manasi Mukherjee** contributed in conceptualization, data curation, investigation, methodology, analysis, writing and reviewing. **Dhriti Banerjee** and **Indu Sharma** contributed to the data on insect abundance across Thar. **Mitali Mukerji** contributed to supervision, writing, reviewing and editing.

